# L-Ascorbic acid restricts *Vibrio cholerae* survival in various growth conditions

**DOI:** 10.1101/2023.04.07.535978

**Authors:** Himanshu Sen, Manpreet Kaur, Saumya Raychaudhuri

## Abstract

Cholera, a deadly diarrheal disease, continues to ravage various parts of the world. It is caused by *Vibrio cholerae*, an important member of the gamma-proteobacteria. Based on certain genetic and phenotypic tests, the organism is classified into two major biotypes, namely classical and El Tor. The El Tor and its variants are majorly responsible for the ongoing seventh pandemic across the globe. Previously, we have shown that cross feeding of glucose metabolic acidic by products of gut commensal can severely affect viability of the biotypes. In this work, we examined the effect of L-ascorbic acid on the survival of *Vibrio cholerae* strains belonging to both biotypes and different serotypes. We observed that L-ascorbic acid is effective in restricting the growth of all strains under various conditions including strains adapted with acid stress.

## Introduction

Cholera, the dreadful diarrheal disease of global importance, is caused by *Vibrio cholerae*. Since its discovery in the 19^th^ century, the organism has been continuously studied worldwide. Over the years of extensive and intensive research has resulted in a wealth of information on the overall biology of the organism with a strong emphasis on the pathogenesis; mode of transmission; survival in host and diverse aquatic environments and epidemiology [1]. Until now, 220 serogroups of *Vibrio cholerae* strains have been reported in the literature of which strains belonging to serogroup O1 and O139 are responsible for cholera epidemic [2, 3]. Other than O1 and O139 serogroup, the strains fall into remaining serogroup are clustered as non-O1, non-O139 which are evidenced to cause local outbreaks [4, 5]. It should be noted that O1 serogroup of *V. cholerae* strains are further classified into two major biotypes, classical and El Tor. As per medical record, the world has experienced seven pandemics of cholera. The first six pandemics are caused by the classical biotype of O1 serogroup strains whereas the El Tor biotype of O1 serogroup strains is responsible for the ongoing seventh pandemic around the world [6]. Although both the classical and El Tor biotypes are closely related, yet some genetic and biochemical differences are still present between the two biotypes including a unique difference in carbohydrate metabolism. It is documented that the viability of classical biotype strains (e.g., O395) is drastically affected in the presence of glucose due to the production of organic acids, whereas the El Tor strains (e.g., N16961) are evolved with the machinery to convert glucose into acetoin, a neutral fermentation end product that promotes the growth of El Tor strains [7]. This evolutionary fitness is believed to be one of the contributing factors on the supremacy of El Tor strains over classical strains in ongoing seventh pandemic which was started way back in 1961 [8]. Interestingly, recent studies have clearly demonstrated that feeding acidic metabolites resulting from the sugar fermentation of gut commensal and probiotics strains is an effective way to restrict the growth and pathogenesis of both classical and El Tor strains under *in vitro* and animal models [9-11]. Since its discovery in 1920, vitamin C has been shown an antibacterial effect against a number of pathogenic bacteria. The molecule also exerts immunomodulatory activity at high concentrations [12]. Recently, sodium salt of vitamin C (sodium ascorbate) has evidenced to act as a quorum sensing inhibitor in *Vibrio campbellii* [13].

To cause cholera, *Vibrio cholerae* must effectively colonize the small intestine. Before reaching the small intestine, pathogen must travel through the acidic barrier of the stomach. To circumvent and survive such hostile environment, *Vibrio cholerae* mounts a powerful acid tolerance response (ATR). The contribution of acid tolerance responses in the infectious life cycle of *Vibrio cholerae* is well recognized [14, 15].

In this work, we examined the effect of L-ascorbic acid on the growth of various strains of *V. cholerae* under various growth conditions. Our data clearly indicated that L-ascorbic acid is effective in restricting the growth of *V. cholerae* strains irrespective of the biotype and serotype examined in the present study. Furthermore, we also demonstrated growth mitigation in acid-adapted *Vibrio cholerae* strains in the presence of L-ascorbic acid.

## Experimental procedures

### Bacterial strains and media

All bacterial strains used for this study are listed in **Table 1**. All *Vibrio cholerae* strains were grown in Lysogeny broth (LB) at 37□ or on solid with agar. All media ingredients were purchased from BD Difco and salts were procured from Sigma-Aldrich. Barritt Reagents A and B were purchased from Himedia.

### Growth assays

Liquid growth assay was set up by diluting the early log phase cultures of *Vibrio cholerae* strains to an OD 600_nm_ of 0.01 in 10 ml of LB supplemented with salts at 37□. Absorbance at 600_nm_ was measured for the stipulated period of time. pH of the cultures was measured.

For viability spotting assays, the liquid cultures were set up as mentioned above. After the stipulated period, the cultures were serially diluted in phosphate buffered saline and spotted on LB agar. Plates were photographed after 16 h of growth at 37□.

### Voges-Proskauer test

Voges-Proskauer (VP) test was performed as described earlier with some modifications [14]. Briefly, *V. cholerae* strains N16961 and C6706 were grown in LB or LB supplemented with 1% glucose (LBG). L-ascorbic acid was added to the LBG media flasks at either 0 h or 4 h of growth. 0.1 ml of each culture was mixed with 0.02 ml of Barritt Reagent A and 0.01 ml of Barritt Reagent B.

### Acid Tolerance response (ATR) and viability assay in the presence of L-ascorbic acid

ATR and viability of *Vibrio cholerae* assay was performed as previously described with slight modifications [14, 17] The overnight cultured *V. cholerae* strains N16961 and C6706 were diluted into fresh LB medium and incubated at 37°C until they reached an OD 600_nm_ of 0.4. The strains were divided into two vials, one having 10% and the other having 90% of the culture. Cells were then washed with LB and then the 10% cells were resuspended in LB pH 7.0 and 90% cells in LB pH 5.7 (pH was adjusted using 1N HCl) and cultured at 37°C for 2 h. Subsequently, the strains cultured at Ph 7.0 or pH 5.7 LB medium were transferred into LB medium at pH 4.5 (adjusted using either HCl or L-ascorbic acid). An aliquot was taken from each culture at the indicated time points and diluted appropriately and spotted on LB agar plates.

## Results and Discussion

### Viability of *Vibrio cholerae* strains is severely compromised in the presence of L-ascorbic acid under various growth conditions

To ascertain the survival of *V. cholerae* strains, *V. cholerae* classical (e.g., O395), El Tor (e.g., C6706), and non-O1, non-O139 strains (e.g., PL91) were subjected to growth assays in Lysogeny broth (LB) containing different concentrations of L-ascorbic acid (L-AA). All strains exhibited severe growth defects in liquid growth assay in the presence of the molecule at concentration of 2.5 mg/ml and 5 mg/ml (**Figure 1A**). The viability spotting also corroborated the same (**Figure 1B**). In addition to *V. cholerae* strains O395, C6706 and PL91, the effect of L-ascorbic acid was examined on more *V. cholerae* strains (e.g. El Tor atypical and non-O1, non-O139, **Table 1**) and observed similar growth inhibition by the molecule (**Supplementary Figure 1**). It should be noted that the addition of L-ascorbic acid caused extreme acidification of the growth medium, which seems responsible for the decimation of *V. cholerae* strains.

**Figure 1.**
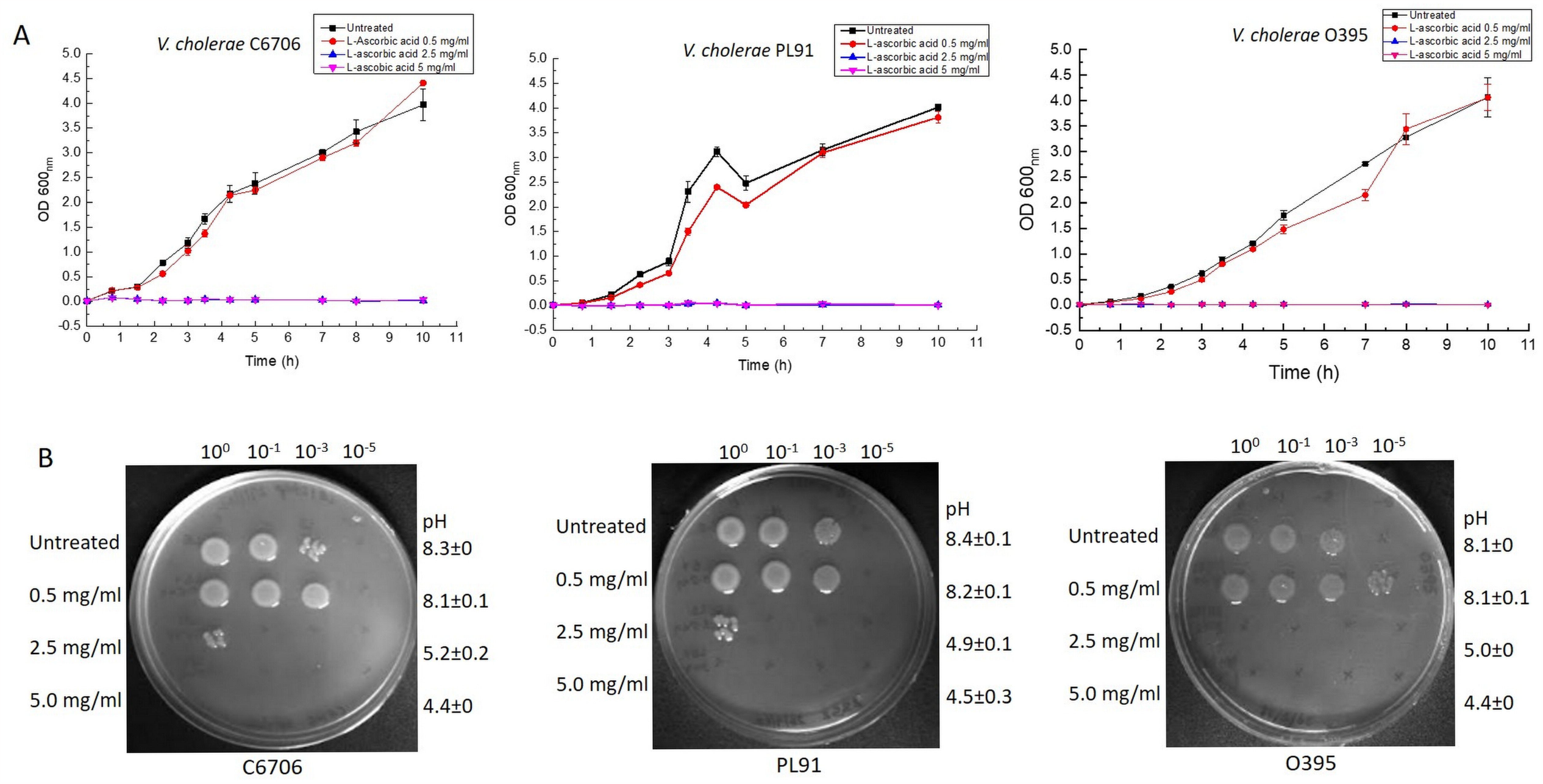
Effect of L-ascorbic acid on *Vibrio cholerae* growth. *Vibrio cholerae* strains C6706, PL91 and O395 were either grown in LB or LB supplemented with increasing concentration of L-ascorbic acid (0.5 mg/ml, 2.5 mg/ml and 5 mg/ml). **(A)** Log phase growth cultures were diluted to a starting OD 600_nm_ 0.01 and OD 600_nm_ was measured at regular intervals. Error bars indicate standard deviation from the mean calculated using the values obtained from biological and technical duplicates of the experiment. **(B)** Cultures grown as in (A) were serially diluted and spotted on solid agar plates. Plates were photographed after 16 h of incubation at 37□. The plate photographs are a representative of the experiment performed in biological and technical duplicates. pH values were measured at the end of the growth assay and standard deviation from the mean was calculated. pH values and their standard deviations were rounded off and reported.

As described in the preceding section, L-ascorbic acid is quite effective in controlling the growth of various *Vibrio cholerae* strains in LB medium. Next, we wanted to examine the effectiveness of the molecule in the presence of ORS salt and glucose. It is noteworthy to mention that some *V. cholerae* strains (e.g., N16961) convert glucose into acetoin (a neutral compound, also serves as an energy source) and gain an advantage on growth in culture medium containing glucose while other *V. cholerae* strains (e.g., O395) produce acidic metabolites as a result of glucose fermentation and encounter a severe challenge to survive under such conditions [7, 9, 10]. Keeping this view in mind, we asked the question whether L-ascorbic acid can tackle such acetoin producing *V. cholerae* strains in glucose containing growth medium. In pursuit of our interest, acetoin positive *V. cholerae* El Tor strains N16961 and C6706 were subjected to grow in LB medium containing 1% glucose and L-ascorbic acid (5 mg/ml) as this concentration was more effective in restricting growth **Figure 1B**) in two ways. In one case, strains were inoculated in LB containing both glucose and L-ascorbic acid and in other case, strains were grown for 4 h in LB containing glucose to produce acetoin (**Supplementary Figure 2**), followed by challenge with L-ascorbic acid and grown for additional 6 h. After 10 h of incubation, the viability was checked by spotting on LB containing streptomycin plate. We observed killing of both the strains under these conditions and pH was found acidic at the end of the incubation (**Figure 2A**). This further indicates that L-ascorbic acid is able to overcome acetoin mediated growth advantage and lowered the pH, thus affecting the survival of these strains.

**Figure 2.**
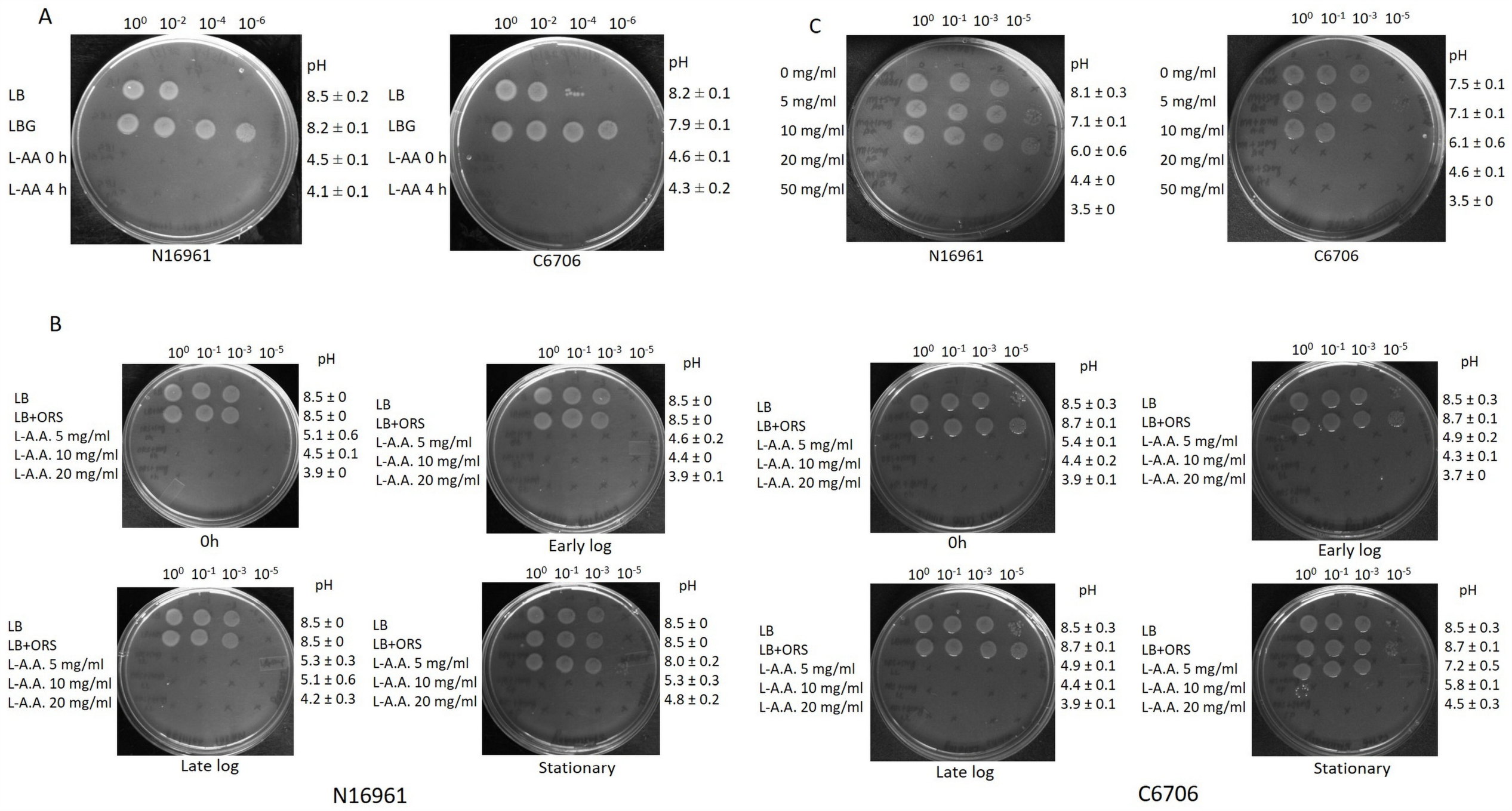
Effect of L-ascorbic acid on *V. cholerae* El Tor strains grown in the presence of glucose and various salts. Log phase growth cultures of *V. cholerae* strains N16961 and C6706 were diluted to a starting OD 600_nm_ 0.01 and **(A)** were grown in either LB or LB + 1% glucose (LBG). L-ascorbic acid (L-AA) (5 mg/ml) was added to the LBG flasks either at the start or after 4 h of growth. **(B)** *V. cholerae* cultures were grown in LB supplemented with the oral rehydration salt mix (ORS). Cultures were challenged with varying concentrations of L-AA (5 mg/ml, 10 mg/ml and 20 mg/ml) at different growth phases [i.e., in the beginning of the assay (0 h), early log phase, late log phase and stationary phase of the cultures]. **(C)** Both El Tor strains of *V. cholerae* were grown in M9 minimal media supplemented with 20 mM casamino acids and increasing concentrations of L-AA (0 mg/ml, 5 mg/ml, 10 mg/ml, 20 mg/ml and 50 mg/ml). For **(A), (B)** and **(C)**, cultures were serially diluted and spotted on solid agar. Plates were photographed after 16 h of incubation at 37□. The plates shown are a representative of the experiment performed in multiple replicates. pH values were measured at the end of each assay and the standard deviation from the mean was calculated. pH values and their standard deviations were rounded off.

To underscore the effect of ORS salts on L-ascorbic acid mediated growth retardation, *V. cholerae* strains N16961 and C6706 were grown in LB containing mixture of salts present in ORS and L-ascorbic acid was added with varying concentrations in various phases of growth. Our viability spotting data clearly demonstrated that increasing concentration of L-AA (20 mg/ml) is effective in complete growth cessation of both strains at stationary phase (**Figure 2B**). Recently, *Vibrio cholerae* is shown to utilize L-ascorbate as energy source. Interestingly, Boyd and colleagues grew *Vibrio cholerae* in M9CAS medium containing glucose or L-AA [20]. On the other hand, we carried out our growth experiments in LB medium. We postulate that M9CAS salt medium neutralizes the acidic condition followed by addition of L-ascorbate, thus promoting growth of the bacteria and the situation further enable *Vibrio cholerae* to utilize L-ascorbate. In order to examine, we grew *Vibrio cholerae* strains in M9CAS medium containing either glucose or L-ascorbic acid at the concentration of 5 mg/ml and 10 mg/ml. No killing was observed at this concentration of L-AA and pH was not acidic either (**Figure 2C**). Therefore, we increased the concentration L-ascorbic acid to 20 mg/ml and 50 mg/ml. Both these concentrations of L-ascorbic acid severely mitigated the growth of *Vibrio cholerae* strains **(Figure 2C**). Collectively, we have demonstrated the efficacy of L-ascorbic acid in restricting growth of different *V. cholerae* strains under various growth conditions.

### L-ascorbic acid mitigates growth of acid adapted Vibrio cholerae strains

As demonstrated, *Vibrio cholerae* strains that are exposed to acid stress can survive at low pH [14]. Therefore, we asked the question whether L-ascorbic acid can affect the growth of acid stress adapted *Vibrio cholerae* strain. To investigate, *V. cholerae* strains C6706 and N16961 were subjected to acid adaptation process as per published protocol [14]. Both unadapted and adapted strains were subsequently exposed to low pH LB media (henceforth LLB pH-4.5) where pH was adjusted either with 1 N HCl (LLB-HCl) or with L-ascorbic acid (LLB-LA). As expected, unadapted strains did not survive in both types LLB media. Interestingly, adapted strain exhibited better survival LLB-HCl medium over LLB-LA medium. In case of LLB-LA, complete elimination of acid adapted *Vibrio cholerae* strains was observed (**Figure 3**). Collectively, our data further reinforced the antibacterial property of L-ascorbic acid that might be exploited to tackle acid adapted variants of *V. cholerae*.

**Figure 3.**
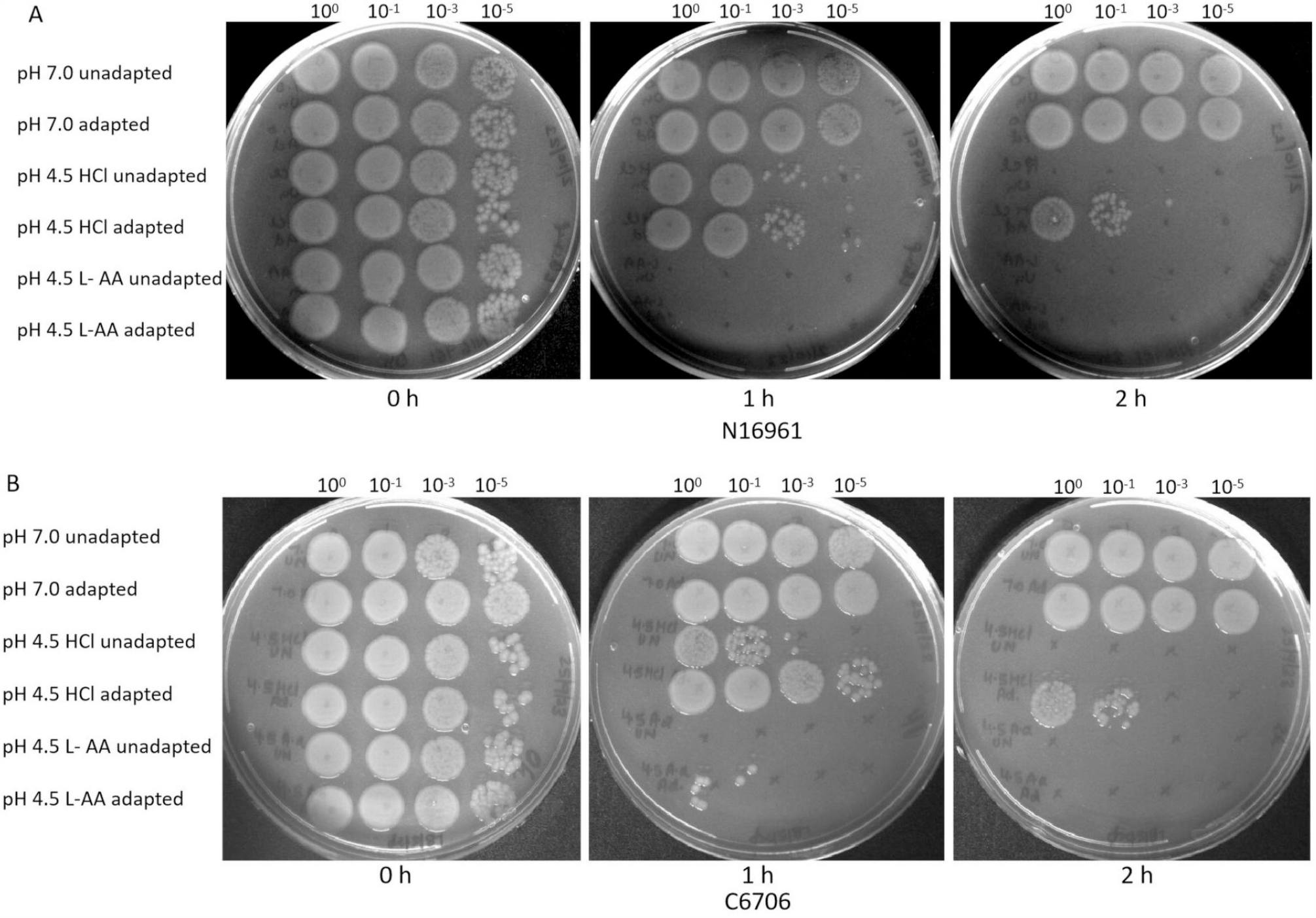
L-Ascorbic acid accelerates the killing of acid adapted *Vibrio cholerae* El Tor strains. Overnight cultures of *V. cholerae* strains (**A**) N16961 and (**B**) C6706 were diluted 1:100 in LB and grown till early log phase. Cultures were then divided in the ratio of 1:9 with one part grown in LB pH 7.0 (unadapted) and nine parts being used for adaptation in LB pH 5.7 (adjusted using HCl) for 2 h. Cultures were then subjected to acid shock in LB at pH 4.5 where pH was adjusted either with HCl or L-ascorbic acid. Serial dilutions were spotted on LB agar plates at time intervals of 0 h, 1 h and 2 h. Plates were photographed after 16 h of incubation at 37□.

The benefits and safety of ascorbic acid are well recognized and established (https://www.chemicalsafetyfacts.org/chemicals/ascorbic-acid/). Recent study also suggested beneficial effect of vitamin C on the gut health [21]. It should be noted that *Vibrio cholerae* invasion causes a tremendous impact on the gut community and some members of gut microbiota play a key role in the cholera recovery phase to restore community structure and function [22, 23]. It would be interesting to examine the effect of combination of ORS with L-ascorbic acid under *in vivo* growth mitigation of *Vibrio cholerae* and restoration of gut microbiota after diarrhoea. Additional studies are necessary to address the issues.

## Authors’ contribution

SRC conceived the idea and designed experiments with HS and MK. HS and MK carried out all experiments. SRC wrote the manuscript. All author gave editorial input and approved the final manuscript.

## Acknowledgements

We gratefully acknowledge Prof. Ron Taylor, Dartmouth Geisel School of Medicine USA, Prof. Richard Kong City University of Hong Kong, Prof. Andrew Camelli, Tuft University and Prof. Roberto Kolter, Harvard University for generously providing strains and plasmids.

## Availability of data and materials

Data sharing not applicable to this article as no datasets were generated or analysed during the current study. Please contact author for data requests.

## Figure Legends

**Table 1. *Vibrio cholerae* strains used in this study**

**Supplementary Figure 1. Growth of *Vibrio cholerae* strains in presence of L-ascorbic acid**. *Vibrio cholerae* strains SC110 and 014-99 were grown in LB supplemented with increasing concentration of L-ascorbic acid (0 mg/ml, 0.5 mg/ml, 2.5 mg/ml and 5 mg/ml). Exponential growth phase cultures of both the strains **(A)** SC110 and **(B)** 014-99 were diluted to a starting OD 600_nm_ 0.01 and grown for 10 h. The serially diluted cultures were spotted on solid agar. The plate photographs are a representative of the experiment performed in biological and technical duplicates. The average and standard deviation of the pH values was calculated from the pH measured at the end of each growth assay. pH values and their standard deviations were rounded off.

**Supplementary Figure 2. Voges–Proskauer (VP) Test of *Vibrio cholerae* El Tor strains to check the production of acetoin**. Log phase growth cultures of *V. cholerae* strains N16961 and C6706 were diluted to a starting OD 600_nm_ 0.01 and were grown in either LB or LB + 1% glucose (LBG). L-AA (5 mg/ml) was added to the LBG flasks either at the start (0 h) or after 4 h of growth. After 10 h, the acetoin production was checked using the VP test for **(A)** *V. cholerae* N16961 and **(B)** *V. cholerae* C6706. The image shown is a representative of the assay performed in biological replicates.

