## Supplementary figures and images for "L-Ascorbic acid restricts *Vibrio cholerae* survival in various growth conditions"

### Supplementary Fig 1

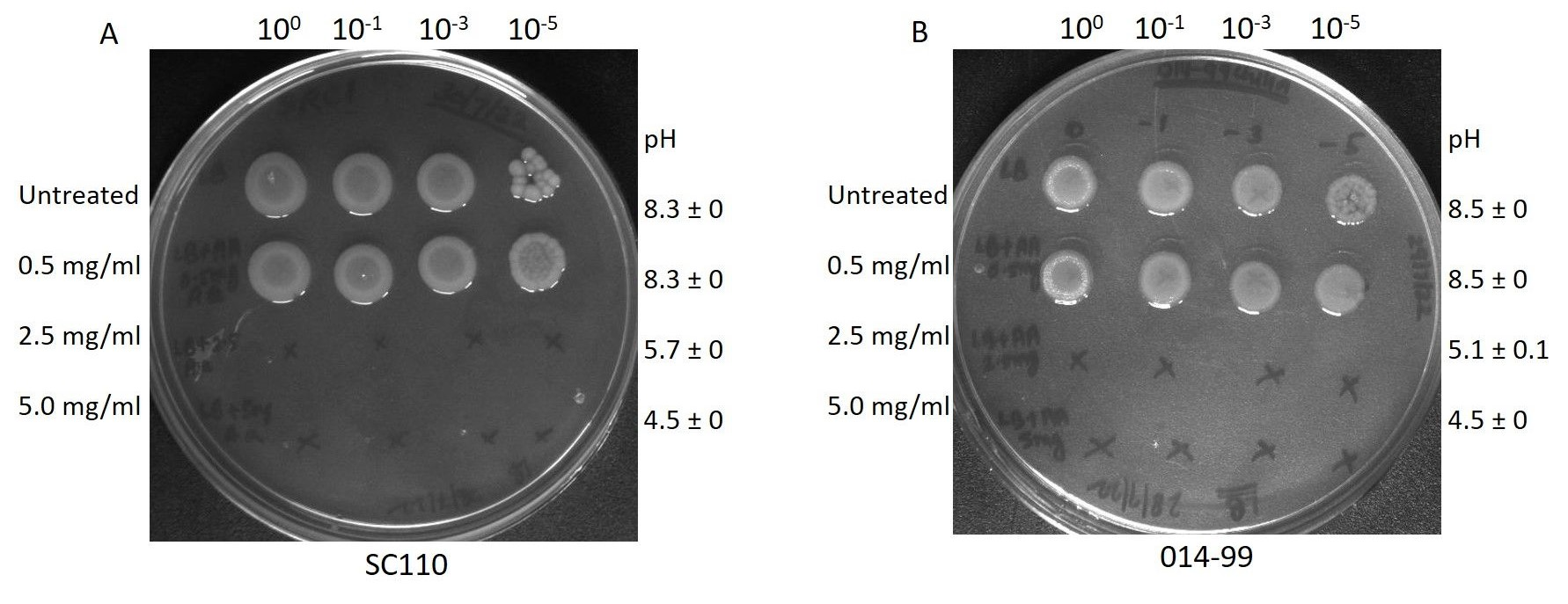

### Supplementary Fig 2

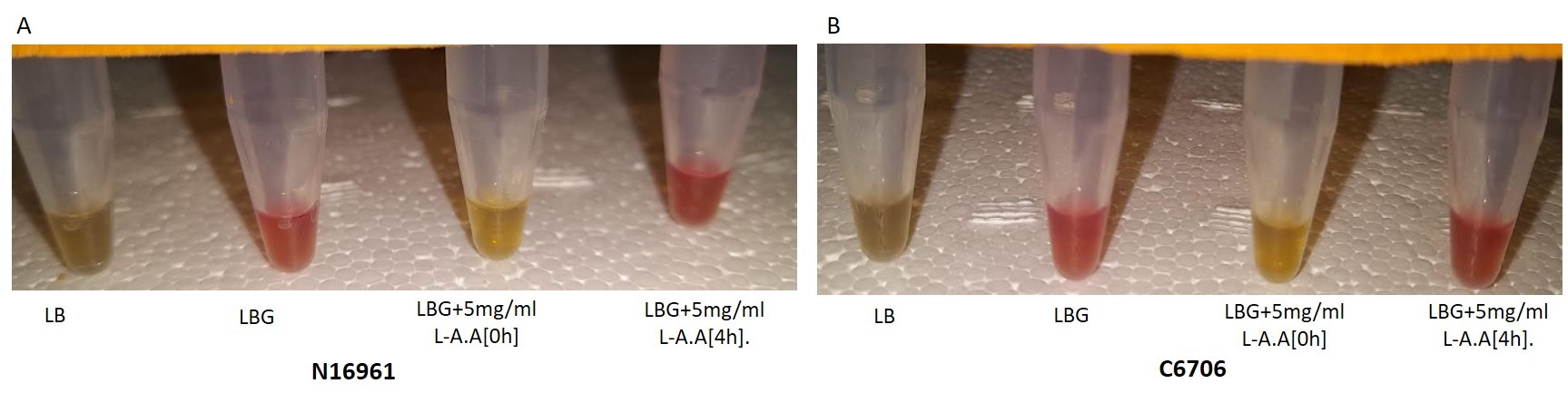
