## Supplementary material for "L-Ascorbic acid restricts *Vibrio cholerae* survival in various growth conditions": Table 1

**Table 1. *Vibrio cholerae* strains used in this study**

| Strain | Genotype | Source |
| --- | --- | --- |
| **C6706** | O1, El Tor, variant, Sm^R^ | Ron Taylor,  Dartmouth Geisel School of Medicine USA |
| **PL91** | Non-O1, non-O139, Serogroup O110, | [18] |
| **O395** | O1, classical | Andrew Camilli,  Tuft University |
| **N16961** | O1, El Tor, Ogawa, Sm^R^ | Andrew Camilli,  Tuft University |
| **SC110** | Non-O1, non-O139 serogroup O34 | [19] |
| **014-99** | El Tor variant | Richard Y.C. Kong,  City University of Hong Kong |
